# Detecting Flavobacterial Fish Pathogens in the Environment Using High-Throughput Community Analysis

**DOI:** 10.1101/2021.06.21.447745

**Authors:** Todd Testerman, Lidia Beka, Emily Ann McClure, Stephen R. Reichley, Stacy King, Timothy J. Welch, Joerg Graf

**Affiliations:** University of Connecticut, Department of Molecular and Cell Biology, Storrs, Connecticut, 06269, USA; Riverence Provisions, Buhl, Idaho, USA; National Center for Cool and Cold Water Aquaculture, Agricultural Research Service/U.S. Department of Agriculture, Kearneysville, West Virginia, USA

**Author notes:** Corresponding author: Joerg Graf, University of Connecticut, Department of Molecular and Cell Biology, Storrs, Connecticut 06269, USA. Geisel School of Medicine, Dartmouth College, Lebanon, NH, USA. College of Veterinary Medicine, Mississippi State University, Starkville, MS, USA.

**Keywords:** aquaculture, pathogen, detection, metagenomics

## Abstract

Diseases caused by the fish pathogens *Flavobacterium columnare* and *Flavobacterium psychrophilum* are major contributors of preventable losses in the aquaculture industry. The persistent and difficult to control infections caused by these bacteria make timely intervention and prophylactic elimination of pathogen reservoirs important measures to combat these disease-causing agents. In the present study, we present two independent assays for detecting these pathogens in a range of environmental samples. Natural water samples were inoculated with *F. columnare* and *F. psychrophilum* cells, and pathogen levels were detected using Illumina MiSeq sequencing and droplet digital PCR. Both detection methods accurately identified pathogen-positive samples and showed good agreement in quantifying each pathogen. Additionally, the real-world application of these approaches was demonstrated using environmental samples collected at a rainbow trout aquaculture facility. These results show that both methods can serve as useful tools for surveillance efforts in aquaculture facilities, where the early detection of these flavobacterial pathogens may direct preventative measures to reduce disease occurrence.

**Importance:** Early detection of a deadly disease outbreak in a population can be the difference between mass fatality or mitigated effects. In the present study, we evaluated and compared two techniques for detecting economically impactful aquaculture pathogens. We demonstrate that one of these techniques, 16S rRNA gene sequencing using Illumina MiSeq technology, provides the ability to accurately detect two fish pathogens, *F. columnare* and *F. psychrophilum*, while simultaneously profiling the native microbial community. The second technique, droplet digital PCR, is commonly used for pathogen detection, and the results obtained using the assays we designed with this method served to validate those obtained using the MiSeq method. These two methods offer distinct advantages. The MiSeq method pairs pathogen detection and microbial community profiling to answer immediate and long-term fish health concerns, while droplet digital PCR method provides fast and highly sensitive detection that is useful for surveillance and rapid clinical responses.

## Introduction

Improving detection methods for bacterial pathogens is a top priority in the global aquaculture industry, as delayed detection and identification of a pathogen can lead to devastating disease outbreaks and high production losses (1). *Flavobacterium columnare* and *Flavobacterium psychrophilum*, the causative agents of columnaris and bacterial cold-water disease (BCWD), respectively, are important threats to the fish farming industry (2). Once introduced into a fish-rearing facility, these pathogens have the potential to rapidly spread and establish biofilms, allowing them to escape biocidal treatments (3, 4). Due to their speed of transmission and long-term survival on surfaces, *F. columnare* and *F. psychrophilum* are immensely difficult to control in fish farms (5, 6) and thus have contributed to major economic losses (7).

Although considerable efforts have been made to control infections caused by flavobacteria over the last fifty years, rainbow trout production losses caused by these pathogens remain high. Studies point to the ability of *Flavobacterium* spp. to survive in a variety of environments, including aquatic and fish-host associated biomes (5, 6). Biofilm formation and the rapid acquisition of antibiotic resistance genes reduce the efficacy of antibiotic therapies for trout. Antibiotics are often ineffective at eliminating pathogens in rearing-facilities and increase the potential for the development of drug-resistant *Flavobacterium* species (8, 9). Genome rearrangement and mutation can also lead to antigen variation in the fish host, reducing the efficacy of new vaccines, potentially explaining why studies report a low reproducibility of results (9–11). The potential of *Flavobacterium* pathogens to linger in fish farms necessitates improved detection methods that allow for early detection and provide insights into potential reservoirs in aquaculture facilities.

Current *F. columnare* and *F. psychrophilum* identification methods range widely depending on the need and capabilities of the laboratory performing the identification (12) and include classical biochemical (13), antigen-based, PCR-based, and DNA array-based techniques (14). The detection of *F. columnare* and *F. psychrophilum* in water and surface samples is further complicated by the presence of multitudinous non-pathogenic *Flavobacterium* species. These environments also commonly contain PCR inhibitors that can reduce the reliability of PCR-based assays (15). Furthermore, *F. psychrophilum* can also be challenging to identify through culture-based approaches because of its slow growth rate and overgrowth by other bacteria (16). These growth characteristics of flavobacteria present a challenge for timely identification, and alternative identification methods are needed.

At aquaculture facilities, similar to natural settings, many areas can harbor pathogens and warrant careful sampling for pathogen detection. For example, after entering a facility with the incoming water, bacterial pathogens can colonize and form biofilms on various surfaces, including the fish raceway walls, containment screens, or baffles (structures commonly used to direct the flow of water downward within raceways). Bacteria in biofilms are more resistant to common biocidal treatment methods, which is a serious problem (17). If the pathogens attach to the surfaces of water pipes or other system components, they can escape disinfection or cleaning attempts and potentially serve as sources of disease recurrence. Many of these issues are of particular concern for flow-through facilities that use untreated spring, ground, or river water. Thus, this specific type of farm could benefit from surveillance at the water inflow. However, even with strategic and frequent sampling, the success of pathogen surveillance and outbreak prevention will be contingent on rapid, sensitive, and specific detection techniques.

Next-generation sequencing (NGS) has been employed in source-tracking pathogens and has the advantage of being able to detect a wide range of pathogens at once, and it is increasingly becoming an affordable tool for use in molecular epidemiology (18, 19). While diagnostic metagenomics is still an unorthodox method in fish disease diagnostics, recent studies on viral detection and persistence in aquatic habitats (20, 21) have demonstrated the potential advantages of this technique. Amplicon deep-sequencing provides an opportunity to detect the presence of lingering etiological agents (20) and may aid in preventing future outbreaks in aquaculture settings.

Marker-gene surveys in particular are common and canonically performed by sequencing a variable region of the 16S rRNA gene, which is ubiquitous and allows for the discernment of bacterial taxa (22). Various approaches are available to analyze 16S rRNA gene sequence data. While traditional approaches use a sequence clustering technique, a more recent approach identifies amplicon sequence variants (ASVs), where the amplified sequence reads are error-corrected, and identical sequences are placed into an ASV. This approach improves pathogen detection sensitivity and is implemented by packages such as QIIME 2 (23) and DADA2 (24). Each ASV is classified by comparing it to a reference dataset. Although the resolution obtained by sequencing a short region of the 16S rRNA gene is not sufficient to identify most taxa at the species level, different variable regions provide greater resolution for some taxa.

Quantitative PCR (qPCR) has long been considered a gold standard for the detection and quantification of disease-causing agents. More recently, droplet digital PCR (ddPCR) has emerged as a promising advancement on qPCR. Due to its partitioning technology and robustness against inhibitory particles, ddPCR has been shown to be an efficient and sensitive approach for detecting low-copy DNA targets and has even begun to replace traditional qPCR in specific applications (25, 26). Additionally, ddPCR results report absolute counts in a sample rather than relative frequencies as are reported with NGS-based methods (27).

In the present study we developed two new assays to detect two pathogenic *Flavobacterium* species, *F. columnare* and *F. psychrophilum*, in water and surface samples by (i) Illumina MiSeq sequencing and (ii) ddPCR. These methods will be useful detection tools for aquaculture facilities. We tested the utility of Illumina MiSeq high-throughput sequencing of the 16S rRNA V4 region to detect these pathogens in inoculated water samples collected from a natural source. Subsequently, these results were independently validated using a ddPCR assay targeting a different housekeeping gene, ATP synthase subunit alpha (*atpA*), which is commonly used as a genetic marker in flavobacterial typing(9). In addition, water and surface samples collected from an aquaculture facility were also tested using both methods, and the results were compared.

## Results

### Differentiation of *Flavobacteria* to the species level using the V4 region of the 16S rRNA gene

A conventional approach for studying microbial communities is to perform high-throughput sequencing of a variable region of the 16S rRNA gene. However, accurate characterization of a polymicrobial sample and detection of specific pathogenic species require that the genetic diversity of the variable region permits species-level classification. Thus, to determine whether the V4 region of the 16S rRNA gene would suffice for species-level identification of fish pathogens of interest, we downloaded the 16S rRNA gene sequences of all 143 available type strains of *Flavobacterium* species from the RDP database to serve as an up-to-date reference gene bank. We trimmed the sequences to the 253 base pairs of the widely used V4 region and constructed a phylogeny. The resulting phylogeny (Figure 1) and the distance matrix (data not shown) identified the closest neighbors of *F. columnare* and *F. psychrophilum* as *F. verecundum* and *F. limicola*, which differed by 4 and 6 base pairs, respectively. Using this approach, the V4 region was observed to provide sufficient resolution to identify *F. columnare* and *F. psychrophilum* to the species level.

**Figure 1:**
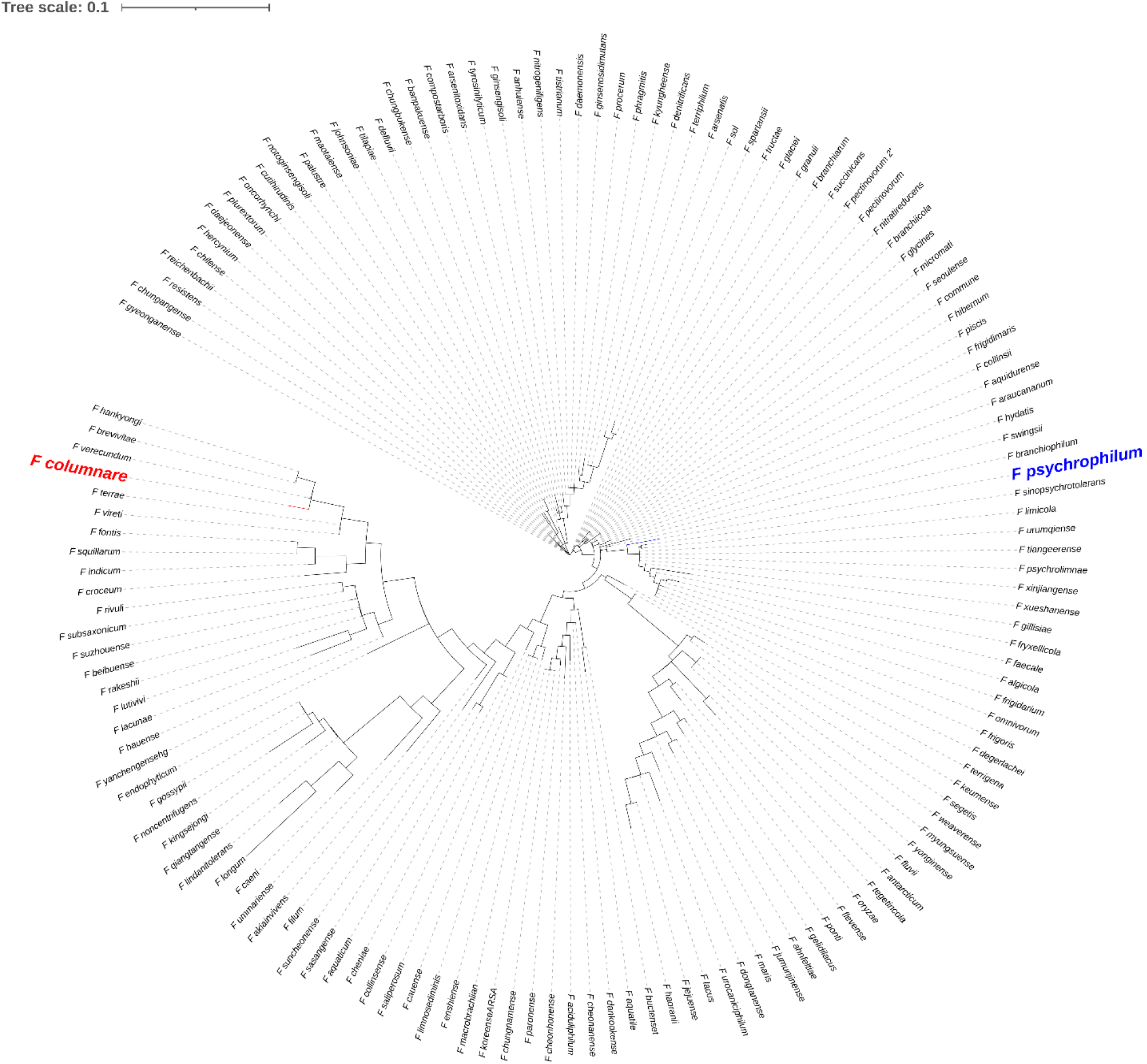
16S rRNA Gene Cladogram for the Genus *Flavobacterium*. Sequences of the 253 base pair V4 region of the 16S rRNA gene from *Flavobacterium* type strains were aligned, and a phylogenetic tree was constructed using Fasttree. *Flavobacterium columnare* (red) is 4 base pairs away from *F. verecundum*, and *F. psychrophilum* (blue) is 6 base pairs away from *F. limicola*.

### MiSeq: Detection of spiked *F. columnare* and *F. psychrophilum* in natural water samples

The ability to detect *F. columnare* and *F. psychrophilum* on an Illumina MiSeq was tested by inoculating and sequencing natural water samples with increasing amounts of these pathogens in three distinct dilution series (*F. columnare* alone, *F. psychrophilum* alone, or their combination). Each sample set was spiked with 10-fold dilutions of pathogen(s) (alone or in combination) over a five order of magnitude range. The relative levels of *F. columnare* and *F. psychrophilum* were normalized using a hemocytometer; however, accurate viable counts were difficult to obtain due to the clumping nature of *Flavobacterium* spp. Two uninoculated samples were used as controls to determine whether native *F. columnare* and/or *F. psychrophilum* could be detected and assess potential filtration system carryover between sample sets. The number of classified *F. columnare* and *F. psychrophilum* was determined using QIIME 2 with a DADA2 plugin for detecting ASVs.

*F. columnare* was detected in all inoculated water samples excluding the lowest-inoculum sample (Figure 2) for both the *F. columnare*-only and co-inoculated sample series. *F. psychrophilum* was detected in all inoculated samples in both dilution series. No sequencing reads were assigned to either *F. columnare* or *F. psychrophilum* in the uninoculated controls. The average read depth of the 17 inoculated samples was 43,853 reads with a standard error of the mean (SEM) of 3,636 reads. A total of 83,143 reads in the entire data set were assigned to *F. columnare* and a total of 38,450 reads were assigned to *F. psychrophilum. F. columnare* was not detected in any samples where it was not expected (samples where *F. columnare* was not added). Only four *F. psychrophilum* reads were recovered from samples where it was not expected (the highest inoculum *F. columnare* sample).

**Figure 2:**
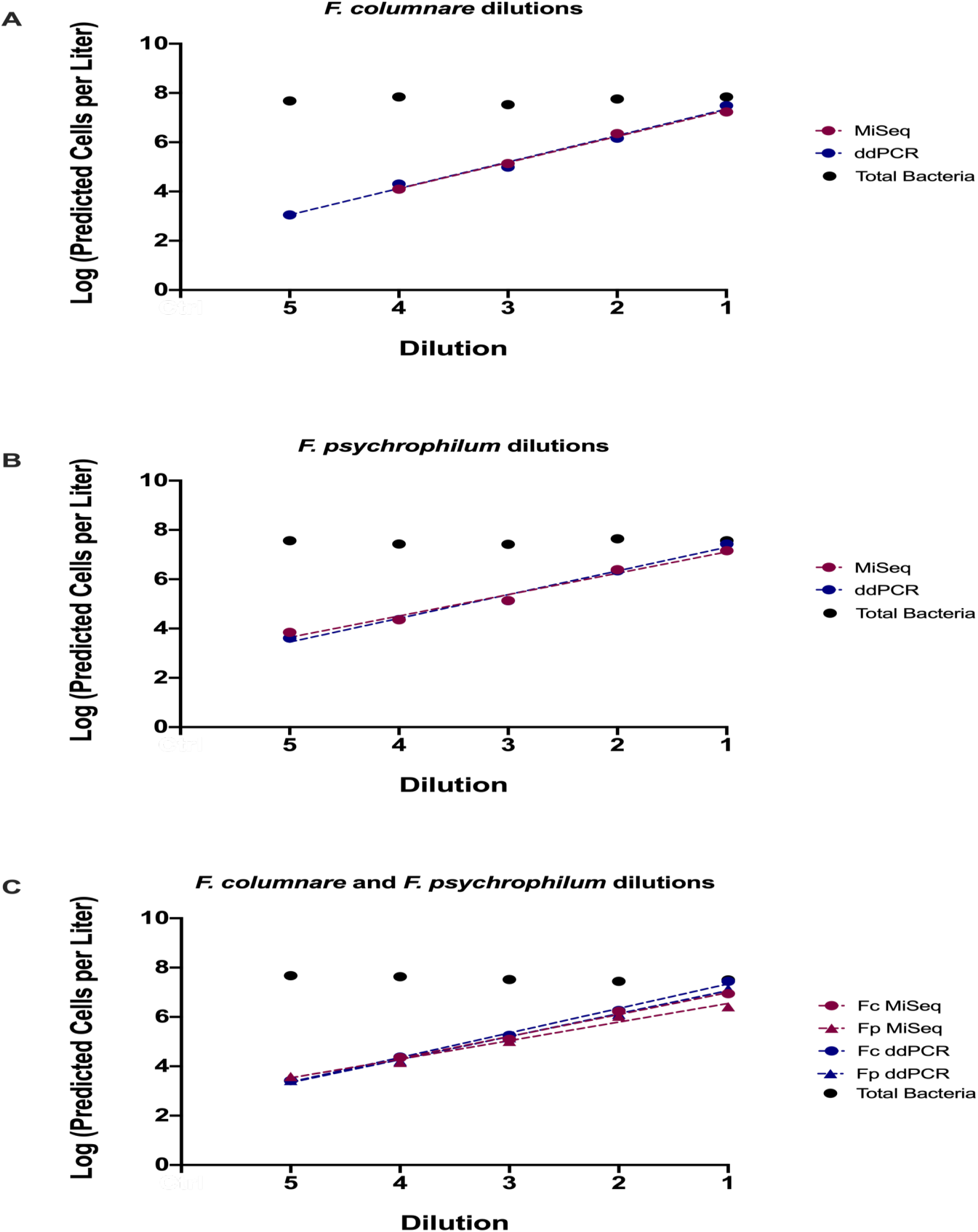
Comparison of MiSeq and ddPCR Assay Results. Estimated cell counts of *F. columnare* (red) and *F. psychrophilum* (blue) are shown for each of the five 10-fold dilutions assayed. Counts are shown on the y-axis as log values, while dilutions are shown on the x-axis and increase from the lowest concentration sample (5) to the highest concentration sample (1). (A) Samples inoculated with *F. columnare* alone, (B) *F. psychrophilum* alone, or (C) *F. columnare*/*F. psychrophilum* combined. The co-inoculated samples (C) show *F. columnare* counts with circles and *F. psychrophilum* counts with triangles. Black circles indicate the total estimated bacterial cell count within the sample calculated using qPCR.

### ddPCR: Detection of spiked *F. columnare* and *F. psychrophilum* in natural water samples

A ddPCR-based approach was developed to independently quantify *F. columnare* and *F. psychrophilum* cells using a single copy housekeeping gene (*atpA*) that has been used previously for species-level discernment in a multi-locus sequence typing scheme for flavobacteria (9). These assays were optimized, validated, and tested for specificity as described in the methods section.

The *atpA*-ddPCR assay was used to determine the number of *F. columnare* and *F. psychrophilum* cells present in the same inoculated water samples evaluated in the Illumina MiSeq detection assay (Figure 2). As expected, the concentrations of both pathogens in each sample series mirrored the 10-fold dilutions over five orders of magnitude using the ddPCR-based assay (Figure 2). We detected ∼1,000 copies of *atpA* per liter of filtered water in the most dilute samples. Since the *F. columnare* and *F. psychrophilum* genomes each harbor one copy of *atpA*, the observed copy number corresponds to 20 genome equivalents/µL of extracted DNA added to the ddPCR reaction mix (total DNA extraction volume was 50 µL).

The ddPCR analysis of samples that were not inoculated allowed us to determine the background levels of *F. columnare* (280 cells/L) and *F. psychrophilum* (12,800 cells/L) in the natural water samples. These counts are much higher than those obtained from the MiSeq experiment, where only a single sample, the highest inoculum *F. columnare* sample, had 3,750 cells/L *F. psychrophilum* detected (four MiSeq reads). This discrepancy could be due to *F. columnare* or *F. psychrophilum* cells being present in the natural water source at levels below the detection limit of the MiSeq method, highlighting the superior sensitivity of ddPCR. Additionally, contamination may have occurred during sample processing or as a result of carry-over from previous filtration operations at levels the MiSeq approach could not easily detect. Lastly, unknown non-target organisms may have cross-reacted with the *F. psychrophilum* assay. To address this discrepancy, the average background level for each method from all non-spiked samples was subtracted from the counts shown in Figure 2.

For ddPCR, a no template control (NTC) as well as two positive controls containing one pathogen and one positive control containing both pathogens were assayed in parallel. The NTC was negative for both pathogens, and only the added target pathogen(s) were detected in the positive controls, indicating that the occurrence of contamination during ddPCR sample preparation is unlikely to explain the background levels of pathogens detected by ddPCR. Droplet counts per reaction ranged from 13,528 to 20,005 droplets with a median droplet count of 18,212. Overall, these data indicate that *F. columnare* and *F. psychrophilum* can be quantified over a wide range of concentrations in natural water samples using ddPCR, but proper controls must be used to ensure accuracy.

### Comparison of MiSeq and ddPCR Results

We were interested in comparing the MiSeq and ddPCR data more directly. As 16S rRNA gene MiSeq data are relative and ddPCR data are absolute, it was necessary to convert absolute numbers while accounting for multiple copies of the 16S rRNA gene in individual bacterial cells. Using a universal 16S qPCR assay, we determined the 16S rRNA gene copy number in each sample (mean = 1.36 × 10^8^ copies, SEM = 1.52 × 10^7^ copies). Additionally, using an estimate of the 16S rRNA gene copy number based on the taxonomic composition of these samples (28), we converted this value to the estimated total number of bacterial cells per sample (mean = 3.6 × 10^7^ cells/mL, SEM = 3.84 × 10^6^ cells/mL; Figure 2, Supplementary Figure 2).

Subsequently, using the absolute concentration data from both the MiSeq and ddPCR assays, we compared the *F. columnare* and *F. psychrophilum* counts in the spiked samples (Supplementary Table 1). Figure 2 displays the line of best fit (dotted lines) produced from a linear regression analysis using log transformed count values. Overall, the results were highly similar between the MiSeq- and ddPCR-generated data and the single and co-inoculation samples. However, for *F. columnare*, the limit of detection differed. In the samples that received the fewest cells, *F. columnare* was not detected using the MiSeq method, whereas ∼1,000 *F. columnare* genome copies were detected with the ddPCR assay (Figure 2a). This difference in detection levels is likely due to the MiSeq producing 10-100 thousand reads and having ∼100,000,000 16S rRNA gene copies in the sample, which may lead to rare sequences being drowned out by more dominant species. Overall, there appeared to be excellent qualitative agreement between the two methods.

On a quantitative level, the estimated cell count values (Figure 2) obtained were highly similar between the two methods. For *F. psychrophilum*, the ddPCR approach yielded higher counts than the MiSeq method for 6 of 10 samples, with an average coefficient of variance (CoV) of 22%. A paired Wilcoxon signed-rank test did not support a significant difference between the methods (p = 0.375). The *F. columnare* counts obtained by ddPCR were higher than those obtained with the MiSeq approach for 6 out 8 samples (only considering samples where signal was detected for both methods), with an average CoV of 25.5%, a difference that was not significant using a paired Wilcoxon signed-rank test (p = 0.25).

### Pathogen Detection in Environmental Samples

We used both the MiSeq and ddPCR methods to evaluate water samples and surface swabs we collected as part of a microbial community survey of an aquaculture facility. Twelve samples were collected from a commercial, spring-fed, rainbow trout farm that had previously experienced periods of BCWD and columnaris disease outbreaks. Table 1 shows the results of the comparison between the two assays for *F. columnare*. In the 6 samples that were positive using the MiSeq method, the ddPCR method also yielded a positive result, while 4 of 6 samples that were negative for *F. columnare* using the MiSeq method yielded extremely low positive results for the ddPCR method. As observed in the previous experiment with spiked samples of lake water, these data also suggest that the MiSeq-based approach of sequencing the V4 region of the 16S rRNA gene is less sensitive than an *atpA*-based ddPCR assay. Alternatively, these results may also suggest that qPCR-based methods are more sensitive to contamination. As none of the aquaculture samples contained *F. psychrophilum* as detected by either method, we could not perform this comparison.

**Table 1:** MiSeq and ddPCR *F. columnare* Detection Results. A positive call using the MiSeq required one or more *F. columnare* reads when normalized to 10,000 reads. A positive call for ddPCR required 40 or more *F. columnare* copies to be detected per 50 µL DNA extract. (+) indicates that less than 2 copies were detected per DNA extract with the ddPCR method or less than 1 read per 10,000 detected with the MiSeq method.

|  | MiSeq (16S-V4) | ddPCR (*atpA*) |
| --- | --- | --- |
| Natural Source Water | (+) | (+) |
| Global Hatchery Outflow Water | + | + |
| Individual Hatchery Pen Outflow Water | + | + |
| Surface Swab from Tailscreen | + | + |
| Adult Raceway Outflow Water 1 | + | + |
| Adult Raceway Outflow Water 2 | + | + |
| Surface Swab from Hatchery Baffle 1 | - | - |
| Surface Swab from Hatchery Baffle 2 | - | (+) |
| Surface Swab from Hatchery Baffle 3 | - | (+) |
| Surface Swab from Hatchery Baffle 4 | - | (+) |
| Gill Swab from Adult Fish | - | (+) |
| Global Hatchery Inflow Water | - | - |

### Microbial Community Analysis of Natural Water Samples

While not the focus of the present study, a high-throughput and multiplexable 16S rRNA gene survey provides useful microbial community composition data as an added benefit. This advantage is particularly notable for the V4 region of the 16S rRNA gene, which serves as an ideal molecular marker for identifying the widest diversity of bacteria (29). Therefore, to further demonstrate the advantages of this approach, we sequenced and analyzed 17 biological replicates of the pelagic community of a freshwater lake (Figure 3, Supplementary Figure 3). As the water was prefiltered through a 2 µm pore sized prefilter, we considered the cells captured on the 0.2 µm Sterivex filter to be planktonic microorganisms.

**Figure 3:**
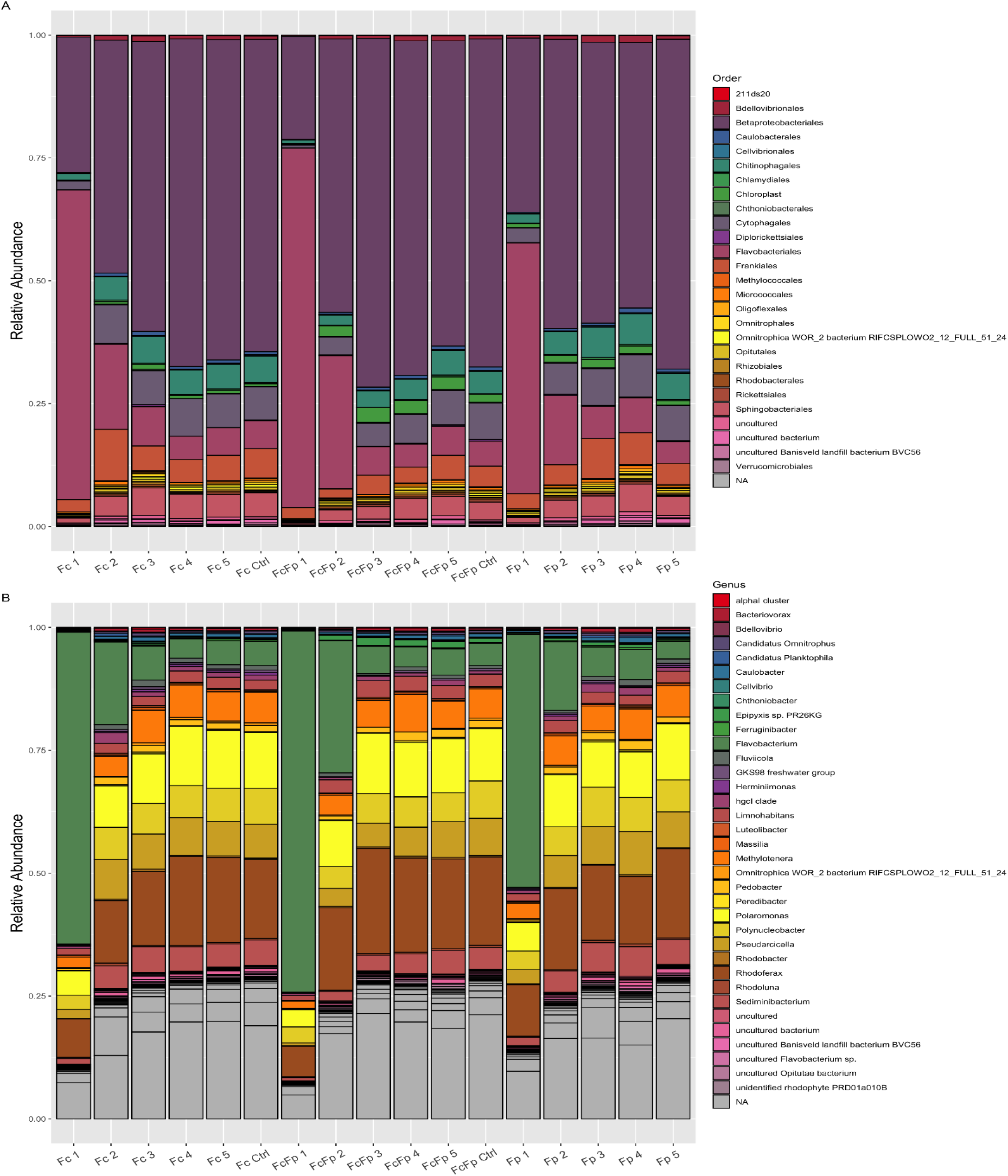
Order and Genus Level Taxonomic Barplots of Water Samples. (A) Order level taxonomic breakdown for each sample. (B) Genus level taxonomic breakdown for each sample. Labels on the x-axis correspond to the dilution series. Barplots are presented as relative abundances. *Flavobacterium* inoculum level decreases from left to right for each series. Fc, Fp and FcFp indicate the addition of *F. columnare* alone, *F. psychrophilum* alone or their combination, respectively. NA – no assignment could be made at this taxonomic level.

At the order level, Betaproteobacteriales dominated the community, accounting for 51.8% of all reads, with constituents present at or above 1% abundance including Flavobacteriales (20.7%), Cytophagales (5.7%), Frankiales (4.9%), Chitinophagales (4.0%), Sphingobacteriales (3.4%), and chloroplast-associated reads (1.2%) (Figure 3A). A visualization of this community with the spiked *Flavobacterium* spp. removed is presented in Supplementary Figure 3A.

At the genus level, 20.0% of reads belonged to the genus *Flavobacterium*, which included the spiked *Flavobacterium* spp. In addition, 14.0, 8.7, 5.5% of the total reads belonged to the genera *Rhodoferax, Polaromonas*, and *Polynucleobacter*, respectively, all of which are members of the family Burkholderiaceae. An additional 14.6% of reads were assigned to the family Burkholderiaceae but could not be confidently assigned to any specific genus. Other genera present at or above 1% included *Pseudarcicella* (5.6%), *Methylotenera* (4.7%), *Sediminibacterium* (3.7%), *Limnohabitans* (2.1%), and *Pedobacter* (1.1%) (Figure 3B). A visualization of this community with the spiked *Flavobacterium* spp. removed is presented in Supplementary Figure 3B.

## Discussion

In the present study, we describe the use of Illumina MiSeq sequencing and ddPCR for the detection of two economically impactful fish pathogens, *F. columnare* and *F. psychrophilum*. We provide direct evidence that MiSeq sequencing of the V4 region of the 16S rRNA gene allows species-level identification of *F. columnare* and *F. psychrophilum*. In addition, we describe a ddPCR assay to detect and quantify these two important fish pathogens, which were detected at similar levels independent of the method used.

Overall, there was good qualitative agreement between the Illumina MiSeq and ddPCR detection of *F. columnare* and *F. psychrophilum*. Statistical testing did not show a significant difference between the quantified levels of *F. psychrophilum* and *F. columnare* using the MiSeq and ddPCR approaches. However, one important difference between the approaches is the limit of detection. Based on the ddPCR quantification data, a detection limit of ∼10^3^ *F. columnare* cells/L was observed for the MiSeq method, while the ddPCR method could detect 10^2^ cells/L. The limit of detection of the MiSeq can be lowered by sequencing to a greater depth, but this would also drive up the overall sequencing costs. A non-targeted NGS-based detection approach requires that the organism of interest be present at a high enough concentration to yield a signal in a complex DNA background (30), which is affected by the sequencing depth. A targeted PCR approach will always outperform a non-specific approach in regard to sensitivity. From a clinical perspective, this difference in the detection limit may be inconsequential, as LD_50_ measurements of *F. columnare* and *F. psychrophilum* are on the scale of 10^8^-10^10^ cells/L for immersion or bath-based infection of rainbow trout (31, 32), well within the detectable range for NGS. Furthermore, due to the slow growth rates of *F. columnare* and *F. psychrophilum*, remediation begun at concentrations of 10^3^ cells/L will likely be effective in keeping concentrations well below the LD_50_

Compared to *F. columnare*, determining the limit of detection for *F. psychrophilum* was more challenging due to an apparent background level of *F. psychrophilum* in the water samples. This may have been caused by the true presence of *F. psychrophilum*, cross-reaction between an off target organism and the primers and probe, or potential carry-over within the filtration system. However, if contamination occurred or *F. psychrophilum* was present in the water, it was not detected by the MiSeq assay, suggesting a cross-reaction with an unknown organism was the likely cause. While the specificity of the primers and probe were bioinformatically verified, the potential for uncharacterized species presents a risk for a false positive signal arising. We observed a baseline of ∼10^4^ *F. psychrophilum* cells in the ddPCR analysis of negative samples. This cross-reactivity was accounted for using a baseline subtraction step for the ddPCR data. By adding a known quantity of target DNA to replicate samples, the presence of cross-reacting DNA can be determined and accounted for in the data analysis.

The benefit of the discovery power added by characterizing the microbial community provided by NGS cannot be overstated. Using this approach, associations between various bacterial taxa and the pathogen of interest can be elucidated, whereas a targeted approach would miss potential hypothesis-generating observations. The potential of other pathogenic bacteria being identified using NGS would also increase the utility of this approach and should be assessed in the future. Compared to ddPCR, a false positive signal arising from NGS is unlikely but possible in the case of primer switching or contamination from wells on the same PCR plate. However, the increased information density produced by this approach increases the odds of correctly parsing out a misidentified sequence. Additionally, it is important to note that because an abundance-filtering step is often performed during pre-processing with NGS datasets (33), low levels of false positive or contaminant reads would be removed as a normal portion of the pipeline. In addition, the inclusion of reagent, negative, and positive controls ensures that contaminants can be more readily detected and removed.

The results obtained for the fish farm samples displayed good agreement between the two methods in detecting the presence/absence of *F. columnare* (Table 1). This result is important, as it was possible to detect this pathogen in both the water and swab samples with the MiSeq. However, the two methods showed disagreement for four surface swab samples in which extremely low levels of *F. columnare* were detected using the ddPCR method but not the MiSeq method. This result may well demonstrate the superior sensitivity of ddPCR in detecting extremely low levels of target sequences. However, this finding may also indicate minor contamination or a PCR artifact leading to a false positive signal. An additional benefit to ddPCR methods is the ability to rapidly process low numbers of samples without increasing the price per sample, while many samples must be assayed at once to decrease the financial input when using the MiSeq.

Although previous studies have demonstrated the potential of NGS in pathogen detection, these investigations typically used shotgun metagenomic sequencing (34, 35) or targeted approaches with genus-specific primers (36). While shotgun approaches can generally provide better resolution, they are significantly more expensive (37), suffer from background or host DNA interfering with detection (34), and require a more complex bioinformatic approach. Targeted NGS methods provide useful information on strain variation within the particular genus of interest but lose the survey capabilities borne from an unbiased approach. In the present study, we demonstrate that specific pathogens with distinct hypervariable regions in the 16S rRNA gene can be reliably and semi-quantitatively detected using a non-targeted NGS method. While not solely intended for use in diagnostic settings, we posit that an NGS-based approach to detect *F. columnare* and *F. psychrophilum* is reliable, has similar processing time compared to culture-based methods (38), and provides additional discovery power that can justify increased cost for a research group.

## Methods

### Phylogeny of *Flavobacterium* type strain 16S rRNA gene sequences

The 16S rRNA gene sequences were downloaded from the Ribosomal Database Project (RDP) database (39) in February 2021. The 253 bp V4 region of the 16S rRNA gene was extracted and aligned using Geneious (version 10.0.6; Biomatters, Auckland, NZ) using a Geneious alignment algorithm. A phylogeny was then constructed in Geneious using Fast Tree (version 2.1.5) (40, 41) with a generalized time reversible model. A Newick file was then exported from Geneious into iTOL (version 6) (42) for additional annotation and export.

### *Flavobacterium* cells and growth conditions

The *F. columnare* ATCC 23463 and *F. psychrophilum* ATCC 49418 type strains were obtained from ATCC (Manassas, VA, USA). Strains were grown on Anacker and Ordall (A&O) agar (43) to reduce clumping (44) and incubated at 22°C for ∼36 h. For liquid cultures, a single colony was grown in A&O broth for 36 h with shaking at 22°C for 36 h.

### Sample filtration

#### Preparation of F. columnare- and F. psychrophilum-spiked water samples

Lake water was collected to ensure that samples would contain a natural freshwater microbial community and have sufficient biomass for downstream processing. Nineteen liters of water were collected on March 29, 2019 from the Mansfield Hollow Lake in Mansfield, CT, USA (41.768568°, -72.174603°). The lake water was prefiltered before use as described in the next section. Then, the cultured bacteria were enumerated with a hemocytometer and added to 1 L aliquots of prefiltered lake water to generate a 10-fold dilution series. To ensure proper mixing, bacterial aliquots were added to 50 mL of sterile water prior to being added to 1 L of the prefiltered lake water.

#### Water filtration

A previously described filtration system (45) was employed in the present study with the following modifications. First, larger particles capable of clogging the system were removed by pumping the lake water sample through a 2 µm pore size prefilter (Sigma Aldrich, St. Louis, MO, USA, Cat # AP2502500) and collected in a sterile vacuum flask. After inoculating the filtrate, the total sample was aseptically mixed and filtered using a 0.2 µm pore size Sterivex filter unit (Fisher Scientific, Waltham, MA, USA, Cat # SVGPL10RC).

One liter of molecular grade water (Fisher Scientific, Cat # BP2819-1) was filtered before the experimental samples and analyzed as a system control to ensure the filtration system was sterile. The samples were filtered in the following order: uninoculated lake water; increasing concentrations of *F. columnare* cells; increasing concentrations of *F. psychrophilum*; uninoculated lake water; and increasing concentrations of 1:1 mixtures of *F. columnare* and *F. psychrophilum*. The filtration system was rinsed with 70% ethanol and molecular grade water (Fisher Scientific, Cat # BP2819-1) between each filtration group before the next sample set. After filtering, the Sterivex filter units were immediately frozen at -20°C.

### DNA isolation

DNA was extracted from Sterivex filters using a Qiagen (formerly MoBio) Sterivex PowerWater DNA Isolation Kit (Qiagen, Hilden, Germany) following the manufacturer’s protocol. DNA was eluted in 50 µL of buffer EB and quantified with a Qubit HS dsDNA Assay Kit (Life Technologies, Carlsbad, CA, USA), with dsDNA concentrations ranging from 4.1 to 11.8 ng/μL.

### 16S rRNA gene amplicon sequencing on an Illumina MiSeq

#### PCR amplification

The V4 hypervariable region of the 16S rRNA gene was amplified in each sample using previously validated primers that contained dual-end adapters for indexing. PCR reactions were set-up in a total volume of 83.4 µL. Reactions contained 3 µL of BSA (New England Biosciences (NEB), Ipswich, MA, USA), 41.7 µL of Phusion 2× Master Mix (NEB), 2.5 µL of 10 µM primer mix (515F forward and 806R reverse rRNA gene V4 primers with Illumina MiSeq adaptors), and 30 ng of sample DNA. Each sample was assayed in triplicate using a Bio-Rad C1000 Touch Thermocycler (Bio-Rad Laboratories Inc., Hercules, CA, USA) with the following parameters: an initial denaturation for 3 min at 94°C followed by 30 cycles of 45 sec at 94°C, 60 sec at 50°C, and 90 sec at 72°C, with a final elongation step of 10 min at 72°C.

#### Library preparation and MiSeq sequencing

Triplicate reactions were pooled and assayed on a QIAxcel (Qiagen) to verify product size and determine DNA concentrations. Samples were then pooled and cleaned using a GeneRead Size Selection Kit (Qiagen, Cat # 180514) before being diluted to approximately 4 nM for loading onto an Illumina MiSeq (Illumina, San Diego, CA, USA). An Illumina 500 cycle MiSeq Reagent Kit v2 (Illumina, Cat # MS-102-2003) was used for paired-end sequencing of all libraries.

### Bioinformatic processing

#### QIIME 2

16S rRNA V4 gene sequence data obtained from an Illumina MiSeq were demultiplexed through BaseSpace (basespace.illumina.com). Paired-end data was then imported into QIIME 2 (version 2019.4; https://docs.qiime2.org/2019.4/), and reads were filtered, denoised, and dereplicated using the DADA2 denoise-paired plugin (24)Sequences were taxonomically classified and compiled using the feature-classifier (46)and taxa (https://github.com/qiime2/q2-taxa) plugins. Taxonomy was assigned using a pre-trained Naïve Bayes classifier based on the Silva 132 99% OTU database (https://www.arb-silva.de/documentation/release-132/), with reads trimmed to only include the region bound by the 515F/806R primer pair. Read counts for taxa of interest were obtained from the csv download function within the barplot visualization provided by QIIME 2. Default QIIME 2 processing parameters were used throughout the workflow except where otherwise noted. All commands and related parameters can be viewed at https://github.com/joerggraflab.

#### ddPCR

Primers and probes were designed using Geneious (version 10.0.6; Biomatters) and are listed in Supplementary Table 2. A 200-bp region of the *atpA* gene was selected for ddPCR analysis using primers and probes unique to *F. columnare* and *F. psychrophilum*. Although a range of primer and probe concentrations were examined for optimum positive/negative droplet separation, the Bio-Rad recommendations of 900 nM primer and 250 nM probe were ultimately used to generate the data shown in the present study. The assay was also optimized using an annealing temperature gradient of 54-59°C. Optimal positive/negative droplet separation occurred at 58°C, with no improvement noted at increased temperatures and a slight decrease in the signal:noise ratio observed at 59°C. An annealing/extension temperature of 58°C was selected for use in all subsequent experiments. *Hae*III was used for restriction digestion of template DNA to decrease droplet rain and, per Bio-Rad recommendations, was included within the reaction mixture after being bioinformatically and experimentally shown not to cut within the target region.

Primer specificity was tested on 21 strains of bacteria including *Flavobacterium* spp. (including the ATCC type strains used in the present study) and other closely related genera. Probe specificity was tested on 9 *F. columnare* strains representing the 4 groups of this species (9). The *F. columnare* assay was shown to be specific for group 1 *F. columnare* (which includes the type strain used in the present study). The *F. psychrophilum* assay was specific for all tested *F. psychrophilum* strains. Neither assay reacted with non-target *Flavobacterium* spp., which included no cross-reactivity between the *F. columnare* and *F. psychrophilum* assays.

Dilution series of *F. columnare* and *F. psychrophilum* in water were used to determine the limit of detection for this method, which was calculated as 1 × 10^−5^ ng of genomic template per reaction with a limit of quantitation of 1 × 10^−4^ ng of genomic template per reaction. Primer specificity was tested by performing ddPCR in the presence of an excess of eukaryotic (fish) or prokaryotic [ZymoBIOMICS Microbial Community DNA Standard (Zymo Research, Irvine, CA, USA)] DNA. At excess DNA concentrations of up to 12 ng/reaction (prokaryotic) or 19 ng/reaction (eukaryotic) (∼1-2 × 10^5^ times more than genomic template), the limit of quantitation was maintained at 1 × 10^−4^ ng genomic template per reaction.

The optimum reaction contained 1× ddPCR SuperMix for Probes (no dUTP) (Bio-Rad), 900 nM forward primer, 900 nM reverse primer, 250 nM each probe, 10 U of *Hae*III (NEB), and >0.1 pg of genomic template DNA. The optimum reaction conditions were as follows:10 min of activation at 95°C activation followed by 40 cycles of (30 sec of denaturation at 94°C and 1 min of annealing/extension at 58°C, with a final incubation at 10 min 98°C for deactivation. Reactions and droplets were prepared according to recommendations outlined by Bio-Rad. Droplets were read and analyzed using a QX200 ddPCR system (Bio-Rad) with the QuantaSoft software package (version 1.7.4.0917).

One-microlter aliquots of DNA extracted from the spiked natural water samples were used to determine total numbers of pathogenic flavobacteria using ddPCR. DNA from samples with high *Flavobacterium* loads were diluted 1:10 to prevent droplet saturation. All reactions were simultaneously assayed in duplex to detect both *F. columnare* and *F. psychrophilum*. For comparison to the MiSeq data, read counts were adjusted by subtracting the average read count from the non-spiked samples (7 samples for *F. columnare* and *F. psychrophilum* each).

#### qPCR

A qPCR assay for 16S rRNA gene quantification was utilized for the present study as previously described (47) (Supplementary Table 2). This assay targets the region of the 16S rRNA gene bound by the 27F and 519R eubacterial primers (48, 49).

qPCR was performed using an EvaGreen (dye-based) assay on a CFX96 Touch Real-Time Thermocycler (Bio-Rad). Reactions were assembled using 2× SsoFast EvaGreen Master Mix (Bio-Rad), 27F and 519R eubacterial 16S primers at 300 nM final concentration, 5 µL of sample or standard DNA, and PCR-grade water to final volume of 20 µL per reaction. Thermocycling conditions were as follows: 98°C for 3 min followed by 40 cycles of 98°C for 10 sec, 55°C for 30 sec with a plate read step directly following each cycle. Data analysis was performed using Bio-Rad CFX Manager (version 3.1). Single threshold quantification cycle (Cq) value determinations were made with a baseline subtracted curve fit. Baseline and threshold values were automatically determined by CFX Manager.

Standards were generated from extracted *F. psychrophilum* genomic DNA. The genome size of the *F. psychrophilum* type strain (2.72 Mb) was retrieved from NCBI and used to calculate the estimated number of copies of the 16S rRNA gene per nanogram of genomic DNA taking into account the gene copy number (6 copies). The standard curve had an efficiency of 111.2% and an R^2^ value of 0.991 with a slope of -3.079 and a y-intercept at 33.262 (Supplementary Figure 1).

All samples (diluted 1:10) fell within the range of the standard curve except two samples; the negative control *F. columnare* sample and *F. psychrophilum* sample with the second-lowest inoculum. These samples failed to amplify at the 1:10 dilution, possibly due to PCR inhibition. For these two cases, a 1:100 dilution was used for quantification, as amplification clearly occurred in these samples. Three NTCs were assayed, and two had detectable Cq values of 26.15 and 30.35. Based on the standard curve, these Cq values correspond to less than 10^2^ bacterial cells in each reaction. This low-level detection is likely due to minor contamination and is not high enough to cause any noticeable change in calculated copies in the actual samples.

The total bacterial cell count was calculated by dividing the 16S rRNA gene count by the estimated median 16S rRNA gene copies per cell in the environment, which was informed by order-level taxonomic profiling using Illumina MiSeq 16S rRNA gene sequencing. The calculation is summarized as follows: the Flavobacteriales read count was multiplied by the V4 copy number of inoculated *Flavobacterium* species plus the Betaproteobacteriales read count multiplied by the median gene copy number (GCN) for this bacterial order in the ribosomal RNA operon copy number database (rrnDB) (50)The resulting value was then divided by the sum of the Betaproteobacteriales and Flavobacteriales read counts. This number varied from sample to sample depending on the proportion of Flavobacteriales to Betaproteobacteriales (the two dominant bacterial orders in these samples). Higher inoculum samples had a higher estimated average 16S rRNA GCN due to *F. psychrophilum* and *F. columnare*, both with higher 16S rRNA GCN than the determined median for the Betaproteobacteriales, making up a larger proportion of the microbial milieu.

#### Controls

Samples that were not inoculated with the target strains were used as additional negative controls, resulting in 7 negative controls for each species. For *F. columnare*, zero reads were detected in the negative control samples. For *F. psychrophilum*, low levels were detected in 1 of 7 negative control samples. Additionally, for MiSeq sequencing, negative and mock community controls were assayed to account for potential contamination and PCR bias issues. Negative controls included PCR and extraction negative controls, neither of which presented a significant number of reads, indicating that contamination was not a major concern at these steps. The mock bacterial community control yielded the expected community composition as specified by Zymo Research.

## Data availability

Raw read data are available in the NCBI SRA database under project ID PRJNA732893.

## Code availability

Source code, including QIIME 2 processing commands and R commands used, is available at https://github.com/joerggraflab/16S-Flavobacterium-Detection.

## Acknowledgments

We thank the UConn Microbial Analysis, Resources, and Services (MARS) facility for performing the sequencing work. We thank Ahmad Hassan and Mariya Riat for helping with the processing of the lake water filtration samples. We thank Dr. Thomas Loch and Dr. Benjamin LaFrentz for providing strains used during the ddPCR assay validation. We thank Dr. Jeremiah Marden for providing feedback and editing the manuscript. This work was funded by the United States Department of Agriculture (USDA) under grant number 8082-32000-006-00-D.

---

**Supplementary Figure 1:**
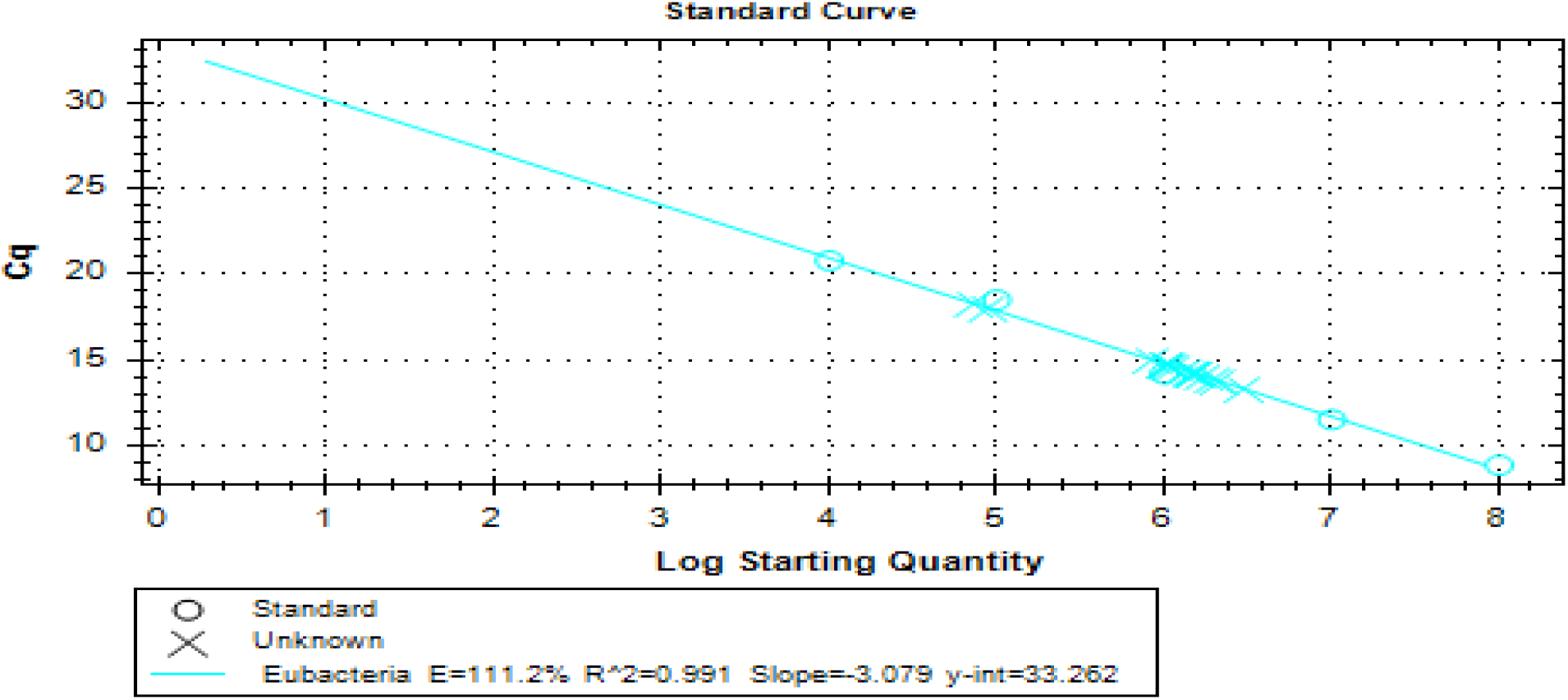
Standard Curve for Eubacterial 16S rRNA Gene Quantification. Circles represent standards and x’s represent samples.

**Supplementary Figure 2:**
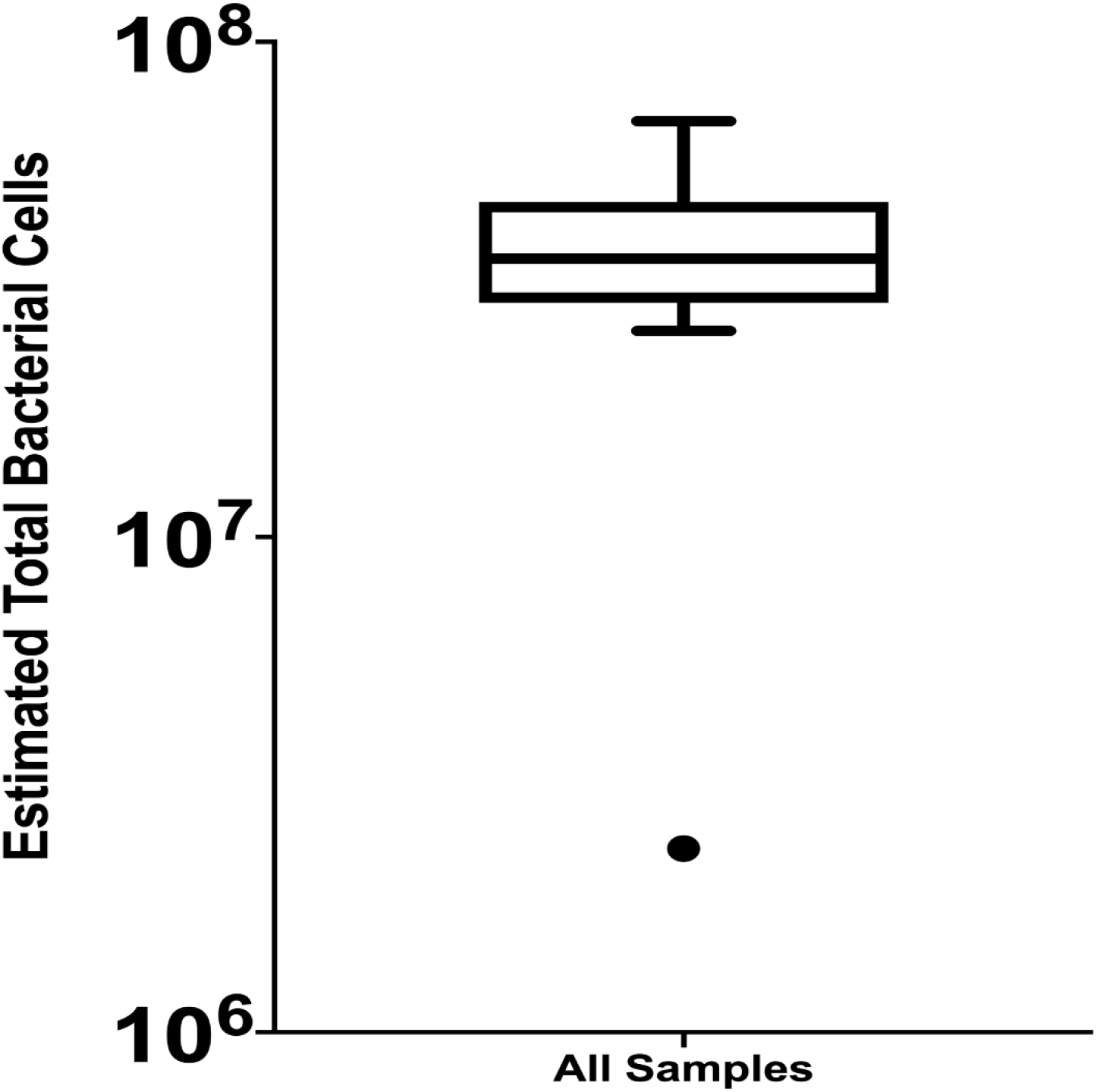
Boxplot of Estimated Total Bacterial Cells as Determined by qPCR. All samples are presented in this boxplot for all dilution series, including negative samples. One outlier is noted that was the negative sample for the dilution series with *F. columnare* alone. The boxplot is presented in Tukey style.

**Supplementary Figure 3:**
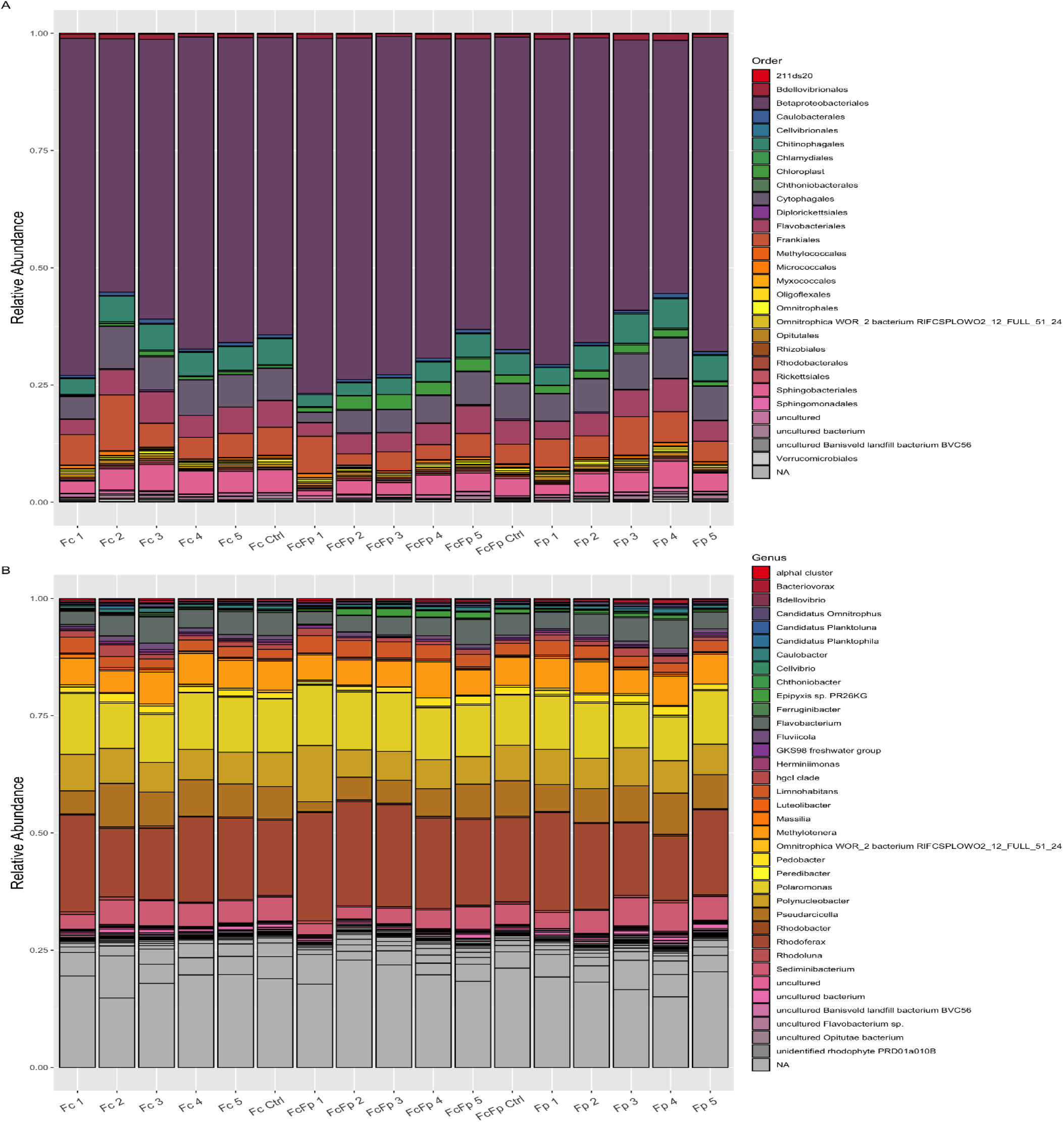
Order and Genus Level Taxonomic Barplots of Water Samples with Spiked Flavobacteria Removed. (A) Order level taxonomic breakdown for each sample excluding reads from *F. columnare* and *F. psychrophilum*. (B) Genus level taxonomic breakdown for each sample excluding reads from *F. columnare* and *F. psychrophilum*. Labels on the x-axis correspond to the dilution series. Barplots are presented as relative abundances. The *Flavobacterium* inoculum level decreases from left to right with each series. Fc – *F. columnare* only, FcFp – Co-inoculation, Fp – *F. psychrophilum* only, NA – No assignment could be made at this taxonomic level.

**Supplementary Table 1:** Calculated *F. columnare* and *F. psychrophilum* Cell Counts for QIIME 2 and ddPCR.

|  | Qiime2 Calculated Fc Cells | ddPCR Calculated Fc Cells<sup>a</sup> | Qiime2 Calculated Fp Cells | ddPCR Calculated Fp Cells<sup>a</sup> |
| --- | --- | --- | --- | --- |
| Fc Ctrl | 0 | 1 | 0 | 1067 |
| Fc 5 | 0 | 1122 | 0 | 0 |
| Fc 4 | 12514 | 20022 | 0 | 0 |
| Fc 3 | 137322 | 97622 | 0 | 2267 |
| Fc 2 | 2238792 | 1460622 | 0 | 0 |
| Fc 1 | 17000414 | 30499622 | 3747 | 8667 |
| FcFp Ctrl | 0 | 0 | 0 | 3267 |
| FcFp 5 | 0 | 2622 | 3833 | 2667 |
| FcFp 4 | 22552 | 23822 | 14408 | 15967 |
| FcFp 3 | 124104 | 180622 | 102359 | 143667 |
| FcFp 2 | 1675120 | 1825622 | 1067381 | 1276667 |
| FcFp 1 | 8784734 | 27799622 | 2608850 | 13637667 |
| Fp 5 | 0 | 422 | 6975 | 4067 |
| Fp 4 | 0 | 0 | 22616 | 23067 |
| Fp 3 | 0 | 0 | 136545 | 135667 |
| Fp 2 | 0 | 0 | 2458902 | 2204667 |
| Fp 1 | 0 | 442 | 14397209 | 26887667 |

a. Cell counts shown for ddPCR are baseline adjusted

**Supplementary Table 2:**
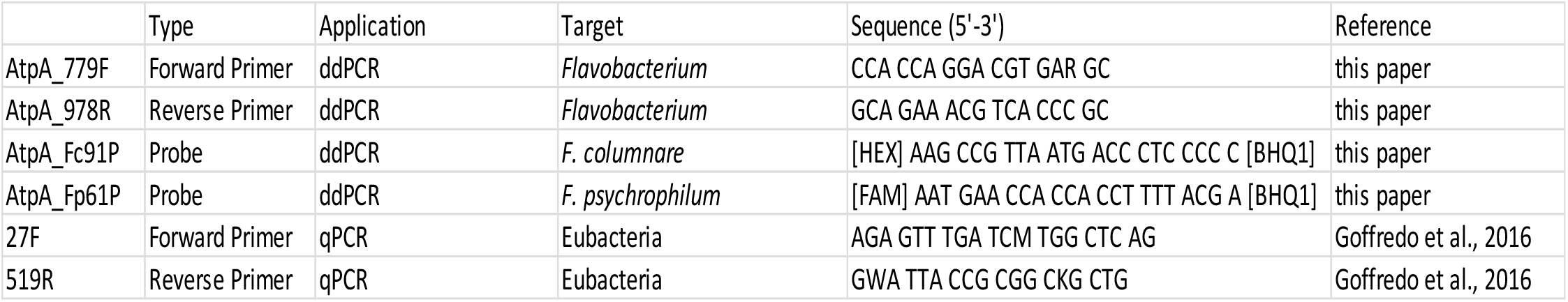
Primer and Probe Sequences Used for ddPCR and qPCR.

